# Cell Painting and machine learning for mechanistic profiling of hepatotoxic botanicals and their constituents

**DOI:** 10.64898/2026.10.05.756246

**Authors:** Guillaume Ollitrault, Constance A. Mitchell, Stephen S. Ferguson, Amy L. Roe, Brandon White, Katherine Titterton, Jessica Ewald

## Abstract

**BACKGROUND:** Botanicals are widely consumed as herbal remedies and plant-derived products, but their complex and variable chemical composition presents major challenges for safety assessment. Although hepatotoxicity has been reported for several botanicals, the constituent compounds responsible and the underlying mechanisms often remain poorly characterised.

**OBJECTIVES:** We aimed to determine whether Cell Painting could identify candidate constituent drivers of botanical bioactivity and generate hypotheses about potential mechanisms of action in primary human hepatocytes.

**METHODS:** We profiled 13 botanical extracts and 85 known constituents across multiple concentrations using Cell Painting alongside assays of metabolic activity and membrane integrity. To interpret morphological phenotypes, we trained machine-learning models using a Cell Painting reference atlas of >1,000 chemicals linked to ToxCast/Tox21 molecular and cellular bioactivity annotations.

**RESULTS:** Cell Painting detected bioactivity in 11/13 botanical extracts and 46/85 constituents, compared with 9/13 extracts and 40/85 constituents in the cell-health assays, and generally detected activity at lower concentrations. For 9/13 extracts, at least one constituent was active at a lower concentration than the corresponding botanical extract, supporting the contribution of a subset of potent constituents to mixture-level responses. Aristolochia and goldenseal were the most potent extracts across Cell Painting and cell-health assays and contained highly active constituents. Reference-atlas models generated specific mechanistic hypotheses, including cytotoxicity-, endocrine-, and genotoxicity-associated responses for several botanicals and constituents.

**CONCLUSION:** These results support Cell Painting combined with machine learning as a sensitive approach for characterising cellular responses to botanicals, prioritising candidate constituent drivers of hepatotoxicity-associated phenotypes, and generating testable hypotheses about potential mechanisms of action.

## Introduction

Botanicals, which comprise a broad range of plant-, algae-, and fungi-derived products, are widely consumed by the general population, often with the aim of supporting health and well-being and, in some cases, for the prevention or treatment of diseases(Bailey et al., 2011; Clarke et al., 2015; Radimer, 2004; Rao et al., 2017). Their use has increased substantially in recent years, with sales of botanical dietary supplements reaching approximately $12 billion in 2022 (Mitchell et al., 2022). Regulatory requirements for botanical safety testing vary worldwide and often depend on the intended use of the product. In several jurisdictions, safety evaluation relies largely on history-of-safe-use information and post-market surveillance rather than systematic pre-market toxicological testing(*Dietary Supplements*, 2005; European Food Safety Authority & EFSA Scientific Committee, 2009). Consequently, many botanical products do not undergo safety assessment to the same extent as other bioactive products such as pharmaceuticals and pesticides prior to human consumption (Bent, 2008). This is particularly relevant for the liver, as it is a key organ for detoxification and xenobiotic metabolism, and is therefore vulnerable to chemically induced injury (Roe et al., 2025). Indeed, botanicals have been associated with liver injury in both clinical reports and experimental studies, and botanical dietary supplements accounted for an increasing proportion of cases documented by the NIH-funded Drug-Induced Liver Injury Network, rising from 7% in 2004–2005 to approximately 20% by 2017(Navarro et al., 2017). Thus, there is a growing need to subject botanicals to chemical safety assessments.

Comprehensive toxicological assessment of botanicals is difficult because they are complex mixtures that can contain hundreds to thousands of phytochemical constituents(Mitchell et al., 2022; Roe et al., 2025). Consequently, even where botanical-associated hepatotoxicity has been reported, the responsible constituents often remain poorly characterised. Scalable new approach methodologies that profile whole botanical extracts alongside their major constituents could help prioritise compounds that may contribute to adverse effects associated with the complete mixture. Prioritised constituents could then be evaluated using established chemical risk-assessment approaches and monitored in future products.

Because botanicals are often consumed with the aim of improving health, an ideal profiling approach would capture both potency and mechanistic information, helping distinguish potentially adverse phenotypes from adaptive or therapeutic responses.

Cell Painting is a high-content imaging approach that is well suited to this type of large-scale potency and mode-of-action characterisation (Seal et al., 2025). The assay uses multiple fluorescent dyes to label key cellular components, including nuclei, mitochondria, endoplasmic reticulum, actin, Golgi apparatus, plasma membrane, and nucleoli (Bray et al., 2016). Cellular representations can then be extracted from these images using conventional feature-based profiling or image embeddings generated by deep learning models such as vision transformers(Oquab et al., 2023; Stirling et al., 2021). Because different biological perturbations can produce distinct morphological signatures, these profiles can be used both to detect bioactivity and to generate hypotheses about potential mechanisms of action.

Previous studies have shown that Cell Painting profiles can partially predict toxicological outcomes including cytotoxicity, mutagenicity, and hepatotoxicity (Cerisier et al., 2025; Lejal et al., 2023; Moshkov et al., 2023; Seal et al., 2021; Way et al., 2021). These properties provide a basis for testing whether botanical phenotypes can be linked to individual constituents and used to generate hypotheses about toxicity-associated mechanisms.

In this study, we profiled 13 botanical extracts and 85 of their constituents across eight concentrations in cultures of primary human hepatocytes using Cell Painting and cell health assays. We used these data to test whether phenotypic profiling could link botanical extract activity to individual constituents that may contribute to hepatotoxicity-associated cellular responses and, through integration with reference Cell Painting and bioactivity datasets, generate hypotheses about potential mechanisms of action. More broadly, this study provides a proof of concept for using high-throughput, high-content screening and machine learning to investigate the cellular effects of complex mixtures and prioritise components for further toxicological evaluation.

## Materials & Methods

### Botanical extracts and constituent compounds

Thirteen (13) botanical extracts including ashwagandha (*Withania somnifera*), usnea *(Usnea spp.*), green tea (*Camellia senensis*), aristolochia (*Aristolochia fangchi* ), kratom (*Mitragyna speciosa*), kava (*Piper methysticum*), milk thistle (*Silybum marianum*), yohimbe (*Pausinystalia johimbe*), comfrey (*Symphytum officinale*), Asian ginseng (*Panax ginseng*), goldenseal (*Hydrastis canadensis*), ephedra (*Ephedra sinica*), and blue cohosh (*Caulophyllum thalictroides*) with 85 of their respective constituents, were supplied by the Botanical Safety Consortium (BSC) for Cell Painting profiling. Details on the extracts, including their source, chemical analyses to support their authentication, and identification and quantification of key constituents can be found at NIEHS’s CEBS website (https://cebs-ext.niehs.nih.gov/cebs/paper/15717), NIH’s database (National Toxicology Program (NTP), 2023), in the supplemental materials, and is described in detail in (Waidyanatha et al., 2024). Botanicals were selected based on existing literature with respect to toxicity or safety, from human (adverse event reporting and clinical trials), animal, or mechanistic studies, and because of their potential hepatotoxicity or lack thereof (described in more detail in (Roe et al., 2025)).

### Cell Painting and cell viability assays

#### Cell culture

384-well microplates (Corning #4518) were prepared by coating with rat tail Collagen I (Thermo Fisher) at 50 µg/mL in 20 mM acetic acid, incubating for one hour, washing twice with sterile H₂O, and air-drying in a sterile cabinet for at least two hours prior to cell seeding. Pooled primary human hepatocytes (PHHs) from BioIVT (Liverpool 5-donor pool) were thawed at 37°C according to the vendor’s protocol, resuspended in INVITROGRO CP medium supplemented with TORPEDO antibiotic mix (Complete CP medium). Cells were resuspended in a total volume of 190 mL and a 30 µL cell suspension was dispensed directly into each dry collagen-coated well at 5,500 cells/well. Plates were centrifuged at 100 × g and transferred immediately to the incubator. Prior to cell seeding, 10 µL of INVITROGRO CP medium was pre-dispensed into each collagen-coated well, and the plates were centrifuged at 216 × g. Subsequently, 20 µL of cell suspension was added to each well, resulting in a seeding density of 6,600 cells/well. Plates were maintained at room temperature for 5 min before transfer to the incubator. Plates were then incubated at 37°C in a humidified 5% CO₂ atmosphere (Cytomat) for 3–5 hours to allow cell adhesion to the collagen substrate.

#### Compound Treatment

Following cell attachment, culture medium was replaced with Complete CP medium containing test substances at twice the final concentration. Extracts were tested in triplicate at eight concentrations, with the upper concentration set by extract-specific solubility limits in DMSO. Constituents were tested in triplicate at eight concentrations ranging from 0.046 to 100 µM: 0.046, 0.14, 0.41, 1, 3.5, 11, 33.5, and 100 µM. The constituent concentration range was selected to approximate levels present within the corresponding botanical extracts based on constituent quantification, expressed as mg/g of extract, and molecular-weight-based conversion to molar units. Plates were returned to the incubator and maintained at 37°C in 5% CO₂ for 44 hours prior to endpoint measurement.

### LDH and MT biochemical assays for cell health assessment

Membrane integrity and mitochondrial activity were evaluated using the CyQUANT LDH Cytotoxicity Assay (Invitrogen) and the RealTime-Glo^TM^ MT Cell Viability Assay (Promega), respectively. For LDH quantification, the assay format differed between the OASIS and BSC datasets. For the OASIS dataset, 2.5 µL of conditioned cell media were transferred to a clear microplate (Greiner 781101) pre-filled with 2.5 µL of medium. Reaction mixture (5 µL) was added and incubated for 30 minutes at ambient temperature, followed by addition of stop solution (5 µL) and absorbance measurement at 530 nm and 720 nm on an SPECTROstar Nano plate reader, per the manufacturer’s instructions. For the BSC dataset, 2.5 µL of conditioned cell medium was transferred to a clear microplate (Greiner 781101) pre-filled with 7.5 µL HBSS. Reaction mixture (10 µL) was added and incubated for 30 min at ambient temperature, followed by addition of 10 µL stop solution, resulting in a total assay volume of 30 µL. Absorbance was measured at 530 nm and 720 nm using an EnVision plate reader. For the MT assay, NanoLuc enzyme and cell viability substrate were diluted in pre-warmed Complete CP medium and applied at 15 µL/well directly to the cell plate, followed by one hour of incubation at 37°C and 5% CO₂ before luminescence reading on the EnVision plate reader. Prior to Cell Painting staining, cell plates were washed twice with HBSS to eliminate residual luminescent substrate that could otherwise interfere with image-based readouts.

### Cell Painting and image acquisition

Fluorescent imaging was performed using the Cell Painting v3 protocol (Cimini et al., 2023) with the PhenoVue Cell Painting JUMP Kit (Revvity, PING21). Following HBSS washing, PhenoVue 641 Mitochondrial Stain was applied at 500 nM in Complete CP medium and incubated for 30 minutes at 37°C and 5% CO₂. Cells were subsequently fixed with 4% paraformaldehyde for 20 minutes at room temperature and washed four times with HBSS. The remaining dyes were then applied simultaneously: PhenoVue Fluor 488 Concanavalin A (5 µg/mL), PhenoVue 512 Nucleic Acid Stain (1 µg/mL), PhenoVue Fluor 555 WGA (1.5 µg/mL), and PhenoVue Fluor 568 Phalloidin (8 nM). The concentration of PhenoVue 512 Nucleic Acid Stain was lowered from the published 3 µg/mL to 1 µg/mL to minimise channel similarity with the adjacent WGA channel; all other steps were performed as described in the original protocol. Images were acquired on an Operetta imaging system (Revvity) with a 40× water objective in non-confocal mode, capturing 15 fields of view per well at full resolution (2,160 × 2,160 pixels, binning = 1). For each of the five fluorescent channels and one brightfield channel, a single z-plane was selected per channel based on maximal Laplacian variance as a measure of focal quality.

### Image and profile processing

#### Feature extraction

Each channel image was pre-processed with instance normalisation at the field level to control for intensity variation. DINOv2 ViT-L/14 model (Oquab et al., 2023), a self-supervised model pretrained on 142 million curated natural images that yields high-dimensional image-level feature was applied independently to each of the six normalised channel images per field of view, generating a 768-dimensional feature vector per channel.

### Profile aggregation and normalisation

Well-level profiles were generated by aggregating field-level features using the median with Pycytominer (Serrano et al., 2025). Profiles were subsequently normalised per plate against DMSO vehicle using the mad_robustize method, followed by feature selection applying variance threshold filtering, removal of columns with missing values, correlation thresholding, and outlier removal in the selection of 4187 features. Concentration-level consensus profiles were derived by aggregating replicate-normalised profiles per compound and per concentration using the modZ method. Compound-level consensus profiles were obtained by taking the median across all concentration-level profiles for each compound.

### Downstream data analysis

#### Botanical extract concentration units

Botanical concentrations were originally prepared and reported in µg/mL, then converted to approximate µM values using an estimated molecular weight for each botanical extract. This suppositional estimated molecular weight was calculated as the average molecular weight of all known constituents associated with each respective botanical (Mauge-Lewis et al., 2025). Because constituent molecular weights vary within each extract, the resulting µM values should be interpreted as approximate estimates.

### Compound similarity

To assess compound similarity between constituents, dimensionality reduction was performed by applying UMAP to the Morgan fingerprints (Rogers & Hahn, 2010) with the Jaccard metric, enabling visualization of compounds in a reduced chemical space. In parallel, hierarchical clustering was conducted based on Jaccard distance with average linkage, allowing the identification of groups of structurally similar constituents.

### Concentration-response analysis

An Interdependence Score (IDS) (Radhakrishnan et al., 2025) was computed for each morphological feature-compound concentration pair to identify features exhibiting a concentration-dependent response for specific compounds. IDS is a computationally efficient method for capturing both linear and nonlinear relationships between two numerical values. We analysed these results to determine whether there were more concentration-dependent features for some channels than others for specific botanicals or constituents using pairwise Wilcoxon signed-rank tests with Bonferroni correction.

The IDS score was also used as a filter to restrict computationally-intensive curve fitting to features with a concentration-response relationship (p-value <0.05 after Benjamini-Hochberg correction). Curve fitting was then performed on selected features and on normalised MT and LDH readouts using fastbmdR (v0.0.0.9) (J. Ewald et al., 2021). Concentrations were log₁₀-transformed prior to fitting. Eight parametric models (Exp2, Exp3, Exp4, Exp5, Poly2, Lin, Power, Hill) were fit to each feature per compound, and the best-fit model was selected based on the lowest residual standard deviation. Fits were excluded if residual standard deviation exceeded three times that of DMSO controls, if the ratio of upper to lower 95% confidence interval bounds exceeded 40, or if the benchmark concentration (BMC) exceeded the highest tested concentration. The BMC was defined as the concentration at which the fitted curve surpassed the 95th percentile of DMSO samples. A compound was considered active if the number of features with a valid BMC exceeded the median number of active features across 100 random permutations, with a minimum threshold of 25 affected features. For active compounds, the point of departure (POD) (also considered as a phenotype-altering concentration) was defined as the 5th percentile of all valid BMCs passing quality filters.Pairwise two-sided Wilcoxon tests were then performed between assays using compounds commonly identified as active, comparing POD values derived from Cell Painting, MT, and LDH assays.

### Clustering of morphological profiles

Clustering of morphological profiles was performed on botanicals and their constituents based on pairwise cosine similarity, using hierarchical clustering with Euclidean distance and Ward’s linkage method.

### Mode-of-action predictions using the OASIS reference dataset

Dinov2 profiles from a Cell Painting dataset that were previously published by the OASIS Consortium obtained downloaded from the Cell Painting Gallery (accession : cpg0037-oasis) (J. D. Ewald et al., 2026; Rouquié et al., 2025; Weisbart et al., 2024). The dataset contains 1085 compounds (https://oasisconsortium.org/oasis-compounds) that were also profiled in primary human hepatocytes using the same experimental protocol and experimental design as the current study. We obtained curated ToxCast assay (Richard et al., 2016) results for OASIS compounds from the GitHub repository associated with the aforementioned study (https://github.com/jessica-ewald/2024_09_09_Axiom_OASIS/tree/main/1_snakemake/inputs/annotations). The OASIS Cell Painting profiles were processed using the same pipeline described above for the botanicals Cell Painting dataset. The two datasets were aligned to account for technical batch effects using the Harmony algorithm (Korsunsky et al., 2019).

XGBoost classification models (Chen & Guestrin, 2016) were trained on compound-level batch corrected morphological profiles from the OASIS dataset to predict binary ToxCast endpoint activity labels. Model development was restricted to endpoints with more than 50 compounds and at least 20 active after label filtering. For each endpoint, models were trained separately on morphological profiles, on cell count features, and on a random baseline, and performance was evaluated using 10 iterations of stratified 5-fold cross-validation with compound-level splits, assessed by mean AUROC and mean AUPRG. AUPRG is a performance metric that evaluates classifier quality using precision–recall gain, reducing the effect of class imbalance and enabling more meaningful comparisons across unbalanced datasets (Flach & Kull, 2015). Only models for which morphological profile-based AUROC and AUPRG were significantly higher than the random baseline across folds and iterations were retained. Retained models were subsequently retrained on their respective full datasets before being applied to predict endpoint activity for BSC botanical extracts and their constituents using the compound-level batch corrected morphological profiles. The ToxCast assays were grouped into biologically relevant main categories using ChatGPT 5.5, based on assay names. The classification process is documented in a shared conversation (Supplementary data 1).

## Results

### Cell Painting detects broad botanical and constituent bioactivity across concentrations

We screened thirteen botanical extracts and eighty-five known phytochemical constituents selected to span a range of reported hepatotoxicity evidence (Figure 1.A, Supplementary Table 1). Two constituents were shared across multiple botanicals: gallic acid, present in green tea and usnea, and epicatechin, present in green tea and kratom. Structural analysis using Morgan fingerprints showed that constituents from the same botanical tended to cluster together, indicating that individual botanicals contained chemically related constituent sets (Figure 1.B). Prior hepatotoxicity evidence varied across the panel: goldenseal, kratom, ephedra, green tea extract, usnea, comfrey, and kava have been associated with hepatotoxicity in humans and/or animal studies, whereas no evidence has been reported for asian ginseng or milk thistle, evidence for yohimbe is inconsistent, and the status of aristolochia, blue cohosh, and ashwagandha remains unclear (Koturbash et al., 2024; Roe et al., 2025).

**Figure 1.**
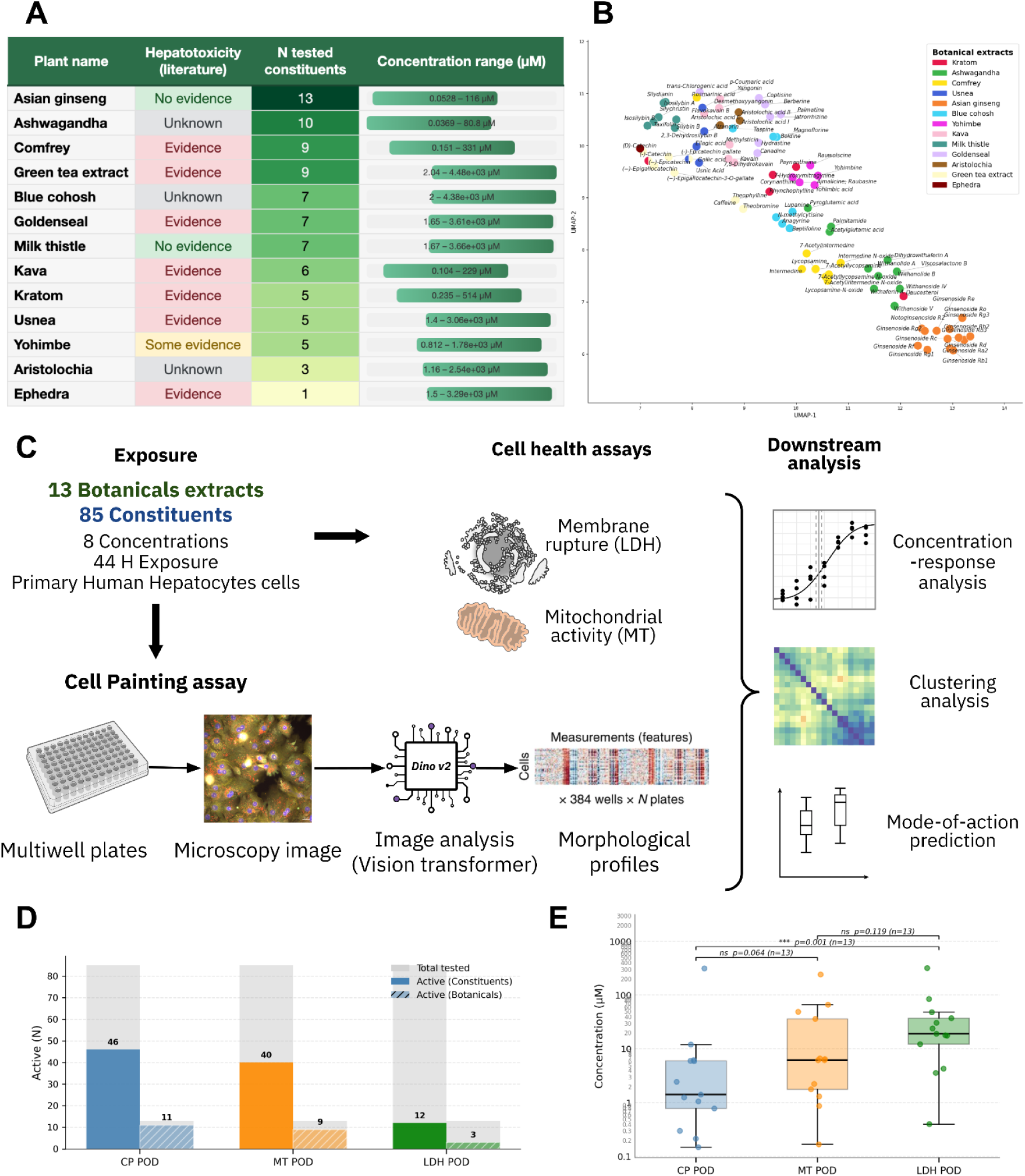
Overview of the evaluated botanicals and their constituent chemical space, and the use of Cell Painting and cell health assays to profile the bioactivity of these botanicals and their constituent compounds. **A.** Summary table of the thirteen botanical extracts examined in this study, indicating the number of constituents tested per botanical and the concentration ranges applied for the extracts, together with prior literature evidence of hepatotoxicity as reported in (Roe et al., 2025) **B.** UMAP projection of the chemical space of botanical constituents, coloured by their parent botanical extract. Dimensionality reduction was performed on Morgan fingerprints using Jaccard distance, illustrating the structural diversity within and between botanical groups. **C.** Schematic overview of the experimental and analytical workflow, including compound treatment and Cell Painting image acquisition through profile processing, concentration-response analysis, reference mapping, and predictive modelling. **D.** Bar plot showing the number of bioactive constituents and botanical extracts identified by Cell Painting, and the MT, LDH cell health assays. **E.** Comparison of points of departure (POD) from MT and LDH Assay with the Cell Painting POD. Only constituents and botanical extracts with measurable activity across all three assays (CP, MT, and LDH) are displayed to avoid bias. Pairwise Wilcoxon signed-rank tests on common compounds are reported.

Primary human hepatocytes were exposed to eight concentrations of each botanical extract and constituent in triplicate for 44 hours, followed by Cell Painting and two complementary cell health assays measuring metabolic activity (MT) and membrane integrity (LDH) (Figure 1.C). Cell Painting profiles were generated using DINOv2 image embeddings. We then used concentration-response modelling to determine whether each treatment was bioactive in each assay and to estimate the concentration at which activity emerged. This enabled us to compare the activity rates and potency of Cell Painting responses with conventional cell health endpoints before examining whether observed botanical phenotypes could be linked to individual constituents and potential mechanisms of action.

Cell Painting detected bioactivity in 56% (48/85) of constituents and 85% (11/13) of botanicals (Figure 1.D, Supplementary Figure 3.B), compared with 49% (42/85) of constituents and 69% (9/13) of botanicals showing activity in either the MT or LDH assays. Among botanicals and constituents active in all three assays (n = 13), activity generally emerged at lower concentrations in Cell Painting than in the MT assay (p = 0.064), with LDH showing the highest PODs (p = 0.001) (Figure 1.E), consistent with previous findings (J. D. Ewald et al., 2026).

We next examined the concordance between previously reported hepatotoxicity and activity across the three assays. Among the eight botanicals with some evidence of hepatotoxicity, seven showed Cell Painting bioactivity and six showed activity in either the MT or LDH assays (Table 1). However, assay activity was not restricted to botanicals with reported hepatotoxicity. Asian ginseng and milk thistle, which have been assessed without evidence of hepatotoxicity, both showed Cell Painting bioactivity, and milk thistle also showed MT activity. Conversely, comfrey, which has been associated with hepatotoxicity, showed no activity in any assay. This was not unexpected, as Cell Painting captures a broad range of cellular responses that may reflect adaptive or intended biological activity as well as adverse effects, whereas endpoints such as LDH leakage more directly indicate cellular injury.

**Table 1.** Bioactivity potency across all measured assays.

| Botanical extracts | Botanical |  |  | Constituents |  |  |  |  |  |
| --- | --- | --- | --- | --- | --- | --- | --- | --- | --- |
|  | CP POD (μM) | MT POD (μM) | LDH POD (μM) | Active CP (n/total) | Median CP POD (μM) | Active MT (n/total) | Median MT POD (μM) | Active LDH (n/total) | Median LDH POD (μM) |
| Goldenseal | 1.41 | 6.46 | 48 | 6/7 (86%) | 2.26 | 7/7 (100%) | 15.45 | 3/7 (43%) | 17.96 |
| Aristolochia | 1.05 | 6.22 | 3.59 | 3/3 (100%) | 6.02 | 3/3 (100%) | 35.97 | 2/3 (67%) | 12.16 |
| Ashwagandha | — | — | — | 4/10 (40%) | 6.18 | 5/10 (50%) | 6.71 | 2/10 (20%) | 27.44 |
| Kratom | 35.71 | 272.9 | — | 4/5 (80%) | 7.24 | 4/5 (80%) | 70.57 | 1/5 (20%) | 56.68 |
| Asian ginseng | 0.22 | — | — | 5/13 (38%) | 7.86 | 4/13 (31%) | 81.28 | 0/13 (0%) | — |
| Blue cohosh | 310.49 | 241.76 | 316.39 | 3/7 (43%) | 9.17 | 3/7 (43%) | 75.8 | 1/7 (14%) | 4.3 |
| Milk thistle | 37.86 | 311.71 | — | 6/7 (86%) | 12.06 | 3/7 (43%) | 66.49 | 2/7 (29%) | 54.81 |
| Usnea | 434.76 | 1990.5<br>2 | — | 3/5 (60%) | 12.06 | 2/5 (40%) | 58.33 | 1/5 (20%) | 84.23 |
| Yohimbe | 74.48 | 209.71 | — | 4/5 (80%) | 13.15 | 4/5 (80%) | 66.71 | 0/5 (0%) | — |
| Comfrey | — | — | — | 4/9 (44%) | 15.21 | 3/9 (33%) | 94.98 | 0/9 (0%) | — |
| Kava | 62.03 | — | — | 3/6 (50%) | 17.19 | 1/6 (17%) | 64.85 | 0/6 (0%) | — |
| Green tea extract | 237.68 | 490.14 | — | 1/9 (11%) | 21.3 | 2/9 (22%) | 58.54 | 1/9 (11%) | 56.68 |
| Ephedra | 318.37 | 1009.3<br>6 | — | 0/1 (0%) | — | 0/1 (0%) | — | 0/1 (0%) | — |
| Median POD / Overall activity rate | 11/13 (85%) | 9/13 (69%) | 3/13 (23%) | 46/85 (54%) | 7.8 | 40/85 (47%) | 66.12 | 12/85 (14%) | 21.48 |

Overall, assay bioactivity showed partial concordance with reported hepatotoxicity but did not clearly distinguish botanicals with and without prior evidence of liver injury. This motivated us to examine whether botanical bioactivity could be linked to specific constituents and whether morphological profiles could provide insight into potential mechanisms of action.

### Botanicals and their constituents induce related responses in cultured primary human hepatocytes

Botanicals are complex mixtures, and screening them alongside their known constituents allowed us to assess whether extract-level bioactivity could be linked to a subset of particularly active compounds. We considered both potency, based on the concentrations at which bioactivity emerged, and phenotypic similarity, based on the Cell Painting profiles produced by each treatment.

#### Botanical bioactivity is often associated with more potent constituent activity

We first compared botanical and constituent potency to determine whether active extracts contained individual constituents capable of producing similar responses at lower concentrations. Of the nine botanicals that affected cell health in the MT or LDH assays, eight had at least one tested constituent that was active in the same assay at a lower concentration than the corresponding botanical extract (Table 1, Figure 2.A). Similarly, among botanicals with measurable Cell Painting activity, nine had at least one constituent with a lower Cell Painting POD than the corresponding extract (Figure 2.B). These results indicate that extract-level activity is frequently accompanied by more potent activity among a subset of constituent compounds.

**Figure 2.**
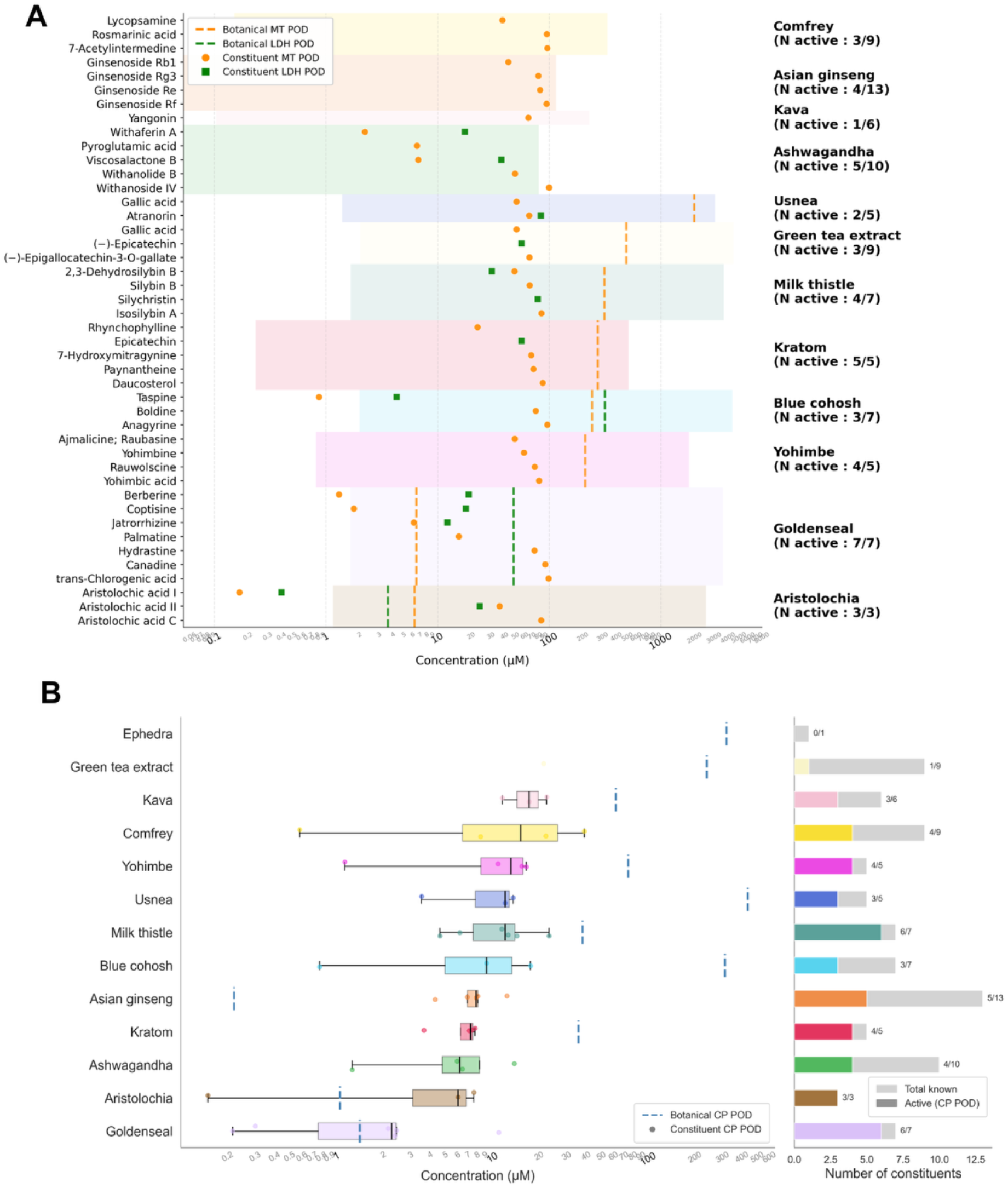
Comparison of constituent-level PODs to botanical PODs. **A.** Comparison of MT and LDH PODs for individual active constituents (points), with the MT and LDH PODs of their respective botanical extracts overlaid as horizontal dashed lines. Background color corresponds to the range of tested concentrations of the botanical. **B.** Boxplots of the constituent CP PODs, grouped by parent botanical extract. Dashed blue vertical lines indicate botanical extract CP POD. Bars in the right panel show the number of constituents active in the Cell Painting assay relative to the total annotated constituents.

Several botanicals showed notable differences between extract- and constituent-level activity. Ephedra showed Cell Painting activity at high concentrations (CP POD = 318 µM), despite none of its tested constituents exhibiting measurable activity; however, several suspected active ephedra constituents are controlled substances (i.e., ephedrine alkaloids) and could not be included in this study. Conversely, ashwagandha and comfrey contained highly active constituents despite no detectable Cell Painting activity from the corresponding extracts within the tested concentration ranges. Withaferin A and intermedine had CP PODs of 1.25 and 0.58 µM, respectively, whereas the maximum tested extract concentrations were 80 and 331 µM. Green tea, usnea, and blue cohosh similarly showed relatively weak extract-level Cell Painting activity despite containing individually active constituents. Asian ginseng showed the opposite pattern, with the lowest extract-level Cell Painting POD (<1 µM) despite limited feature activity, no detectable cell health effects, and constituents that were consistently less potent (CP PODs 4.29–12.41 µM). By contrast, milk thistle and kava showed closer agreement between extract and constituent activity, with extract CP PODs falling within the interquartile range of their constituent PODs.

Aristolochia and goldenseal were the most potent botanical extracts across both Cell Painting and cell health assays. Both showed cell health activity below 10 µM and Cell Painting activity below 2 µM. For aristolochia, aristolochic acid I was also among the most potent constituents across both assay types, consistent with it being a plausible contributor to extract bioactivity. Goldenseal similarly contained several highly potent constituents, including berberine, coptisine, jatrorrhizine, hydrastine, and palmatine, with berberine ranking as the second most potent constituent overall. Together, these results show that highly active botanical extracts can often be linked to a small subset of potent constituents, while also revealing cases in which extract-level activity cannot be readily inferred from the tested constituents. Cell Painting reveals related phenotypic responses across botanicals and constituents

We next asked whether botanical extracts and constituents produced similar morphological phenotypes, providing additional information beyond their relative potency. Cell Painting profiles were aggregated at the treatment level and compared using hierarchical clustering of profile similarity, enabling phenotypic relationships among botanicals and constituents to be examined across the dataset (Figure 3.A). The most prominent cluster contained botanicals and constituents that were also active in the cell health assays (cluster 1, Figure 3.A), consistent with cytotoxicity and associated changes in cell density producing strong morphological signatures (Seal et al., 2021). In contrast, clusters 2 and 3 contained treatments that were inactive in the cell health assays, or active only at relatively high concentrations, despite showing Cell Painting bioactivity. Cluster 2 included Asian ginseng and its constituent ginsenoside Rg2, while cluster 3 included the kava constituents flavokavain B and desmethoxyyangonin. These clusters therefore captured morphological responses that were distinct from the dominant cell-injury-associated phenotype and may reflect additional biological processes. Together, these results indicate that Cell Painting can distinguish both broad cell-health-associated responses and more specific phenotypic similarities among botanicals and their constituents. Such similarities provide a basis for generating hypotheses about shared cellular responses that can subsequently be investigated using more targeted mechanistic analyses.

**Figure 3.**
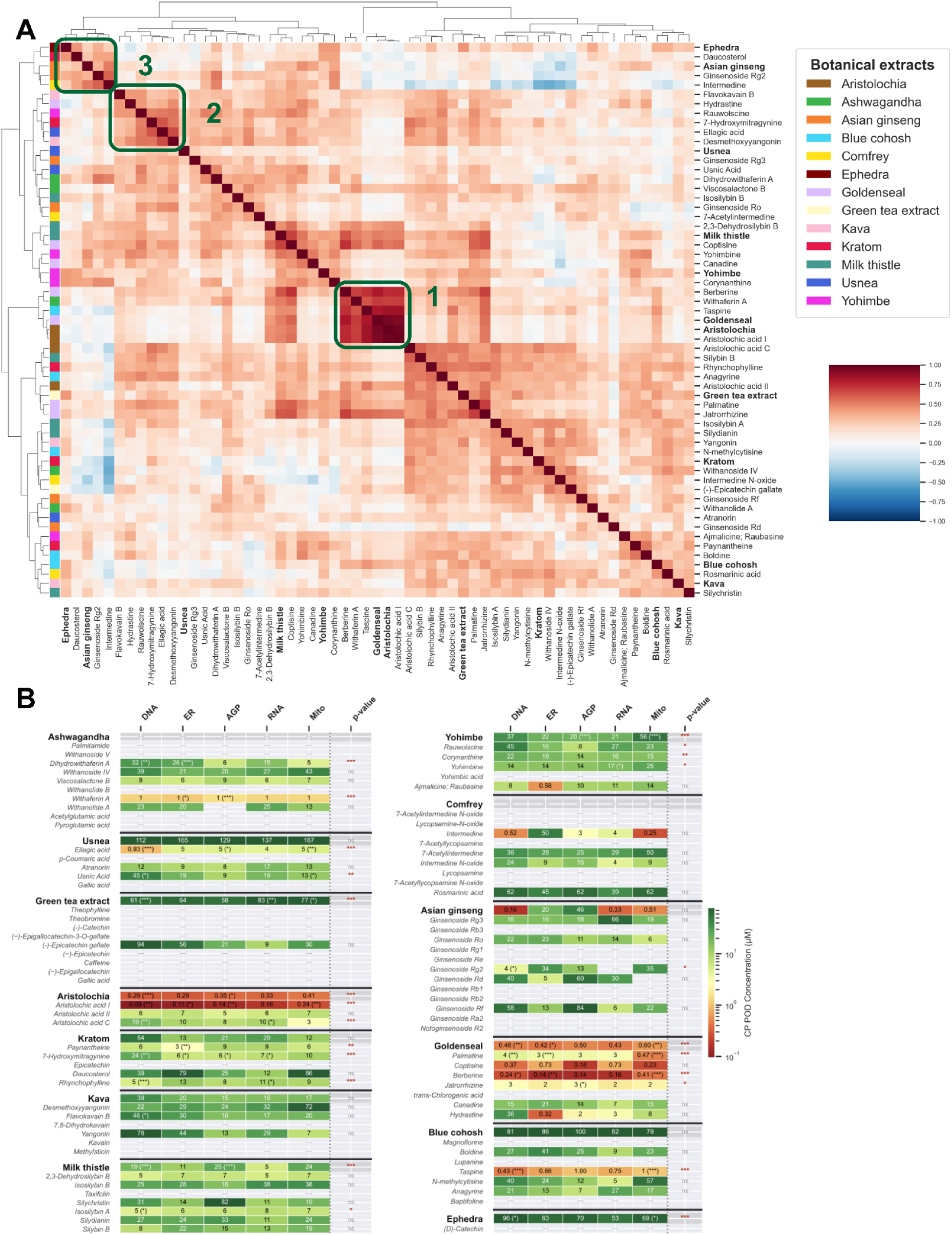
Cell Painting reveals phenotypic similarities and compartment-specific responses across botanicals and constituents. **A.** Clustermap of morphological profiles aggregated at the compound level of active constituents and botanical extracts. Profiles were clustered using euclidean distance on the cosine similarities using the ward linkage hierarchical clustering. **B.** Table of channel specific CP PODs with marked significant POD comparison across channels.

### Compartment-specific Cell Painting responses provide additional mechanistic clues

Because individual Cell Painting channels capture different cellular compartments, we next examined whether botanical and constituent responses differed across channels. Across all substances, the mitochondrial channel contained the greatest number of features significantly associated with concentration, whereas the DNA channel contained the fewest (Supplementary Figure 5.A). Comparison of channel-specific PODs identified 23 substances with significant differences in potency across cellular compartments (Figure 3.B). Several of these compartment-specific responses were consistent with known biological activities. For example, aristolochic acid I produced a DNA-channel POD of 0.09 µM, the lowest channel-specific POD observed across all constituents. This pronounced DNA-associated response is consistent with the well-established genotoxic and carcinogenic properties of aristolochic acid I (Das et al., 2022; Roe et al., 2025). More broadly, these results show that separating Cell Painting responses by cellular compartment can reveal phenotypic signals that may be obscured in whole-profile summaries and provide additional clues for interpreting potential mechanisms of action.

### Supervised machine learning generates specific mode-of-action hypotheses for botanicals and their constituents

Unsupervised clustering revealed groups of botanicals and constituents with similar phenotypic responses that may reflect related modes of action (Figure 3.A). However, because Cell Painting features have limited direct mechanistic interpretability, unsupervised similarity alone cannot readily assign specific biological processes to these phenotypes. One approach is to learn morphological signatures associated with known cellular responses from a reference set of well-characterised compounds and then apply these models to less-characterised treatments.

#### Botanical profiles can be aligned with a hepatotoxicity reference atlas

Our botanical dataset contained too few compounds with established modes of action to train robust supervised models directly. We therefore aligned the profiles to a larger reference dataset generated by the Omics for Assessing Signatures for Integrated Safety (OASIS) Consortium (J. D. Ewald et al., 2026; Rouquié et al., 2025). The OASIS dataset contains 1,085 relatively well-characterised compounds profiled across eight concentrations (0.01–100 µM) using a nearly identical experimental protocol, although the datasets were generated in different technical batches approximately eight months apart. After Harmony batch correction (Korsunsky et al., 2019), OASIS and botanical profiles occupied overlapping phenotypic space in UMAP representations at both the replicate and compound-consensus levels, including for DMSO controls (Figure 4.A, Supplementary Figures 6.A and 9). Exposures producing strong cell health effects also occupied similar regions across the two datasets (Supplementary Figure 4.A), indicating that related phenotypic responses were represented similarly after alignment. Together, these analyses supported the application of models trained on the OASIS reference dataset to the botanical profiles.

**Figure 4.**
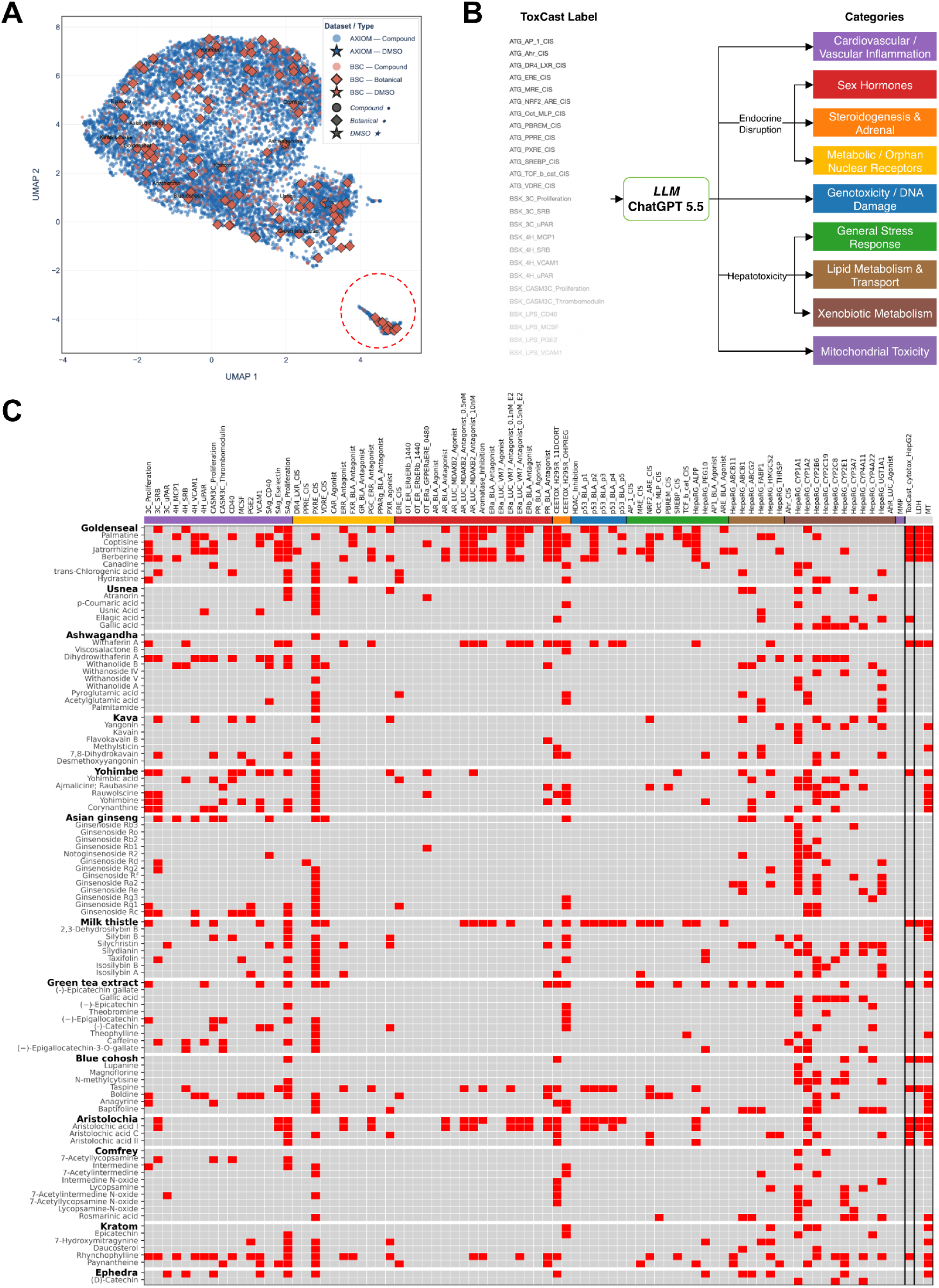
Prediction of botanical extracts and their constituents modes-of-action using the OASIS reference compounds set. **A**. UMAP of the profiles aggregated at the concentration level. The dashed red circle corresponds to compound-concentration pairs highly active in cell health assays. **B**. Categories of ToxCast assays used to make predictions of botanical mode-of-action. **C.** Heatmap showing model predictions for botanical extracts and their individual constituents across the different assays. Each point represents the predicted activity of a given extract or compound in a specific assay. Red points indicate an active prediction, whereas grey points indicate an inactive one.

### Machine learning predicts molecular and cellular responses from Cell Painting profiles

Most compounds in the OASIS reference dataset were also profiled by the ToxCast/Tox21 screening program (Richard et al., 2016), providing molecular and cellular activity labels for supervised learning (J. D. Ewald et al., 2026). For each assay represented by at least 50 compounds and 20 active hits, we trained an XGBoost classifier to predict assay activity from OASIS Cell Painting profiles (Supplementary Figure 7). Of 139 classifiers, 83 (60%) performed significantly better than a random baseline (Supplementary Figure 10). These endpoints spanned processes including endocrine signalling, xenobiotic metabolism, inflammation, genotoxicity, mitochondrial dysfunction, and cytotoxicity (Figure 4.B, Supplementary Table 2). The OASIS dataset also included MT and LDH measurements collected alongside Cell Painting. Models trained to predict these endpoints showed the highest performance among the classifiers tested (MT AUROC = 0.87; LDH AUROC = 0.93). We retained these models together with the 83 significantly predictive ToxCast/Tox21 classifiers, yielding 85 classifiers that were subsequently applied to botanical and constituent profiles to predict activity across the corresponding endpoints (Figure 4.C).

Before interpreting the predicted modes of action, we assessed how well the models transferred to the botanicals dataset using two complementary analyses. First, since our study also measured MT and LDH assays, we could directly evaluate their predictive performance. We found that the Cell Painting models predicted the MT and LDH assays well beyond the random baseline for the botanical extracts and constituents (AUROC = 0.76 for MT and 0.75 for LDH). Second, we compared the biological coherence of OASIS assay predictions to our botanical assay predictions, using the 418 OASIS compounds that had no ground truth ToxCast annotations. We observed that related biological endpoints tended to be predicted as active for the same compounds to a similar extent for both datasets (Supplementary Figure 8A,B). Together, these results suggest that the trained classifiers make plausible predictions when applied to the botanicals dataset.

We next examined the prevalence of predicted activity across the reference endpoints. Among the most frequently predicted responses were activation of the pregnane X receptor (ATG_PXRE_CIS; 60%, 59/98), effects on cell proliferation (BSK_SAg_Proliferation; 44%, 43/98), induction of CYP1A1 in HepaRG cells (LTEA_HepaRG_CYP1A1; 43%, 42/98), and altered mitochondrial activity measured by MT (26%, 26/98). These predictions suggest that botanical-induced morphological responses frequently capture processes related to xenobiotic metabolism, proliferation, and cell health.

At the botanical level, milk thistle showed the highest proportion of predicted active endpoints (32.9%), followed by goldenseal (31.8%) and aristolochia (28.2%). Goldenseal and aristolochia were also among the most bioactive botanicals experimentally, while milk thistle showed measurable activity at higher concentrations. In contrast, ashwagandha (1.2%) and comfrey (2.4%), which showed little or no experimental extract-level activity, had few predicted active endpoints. Kratom showed relatively few predicted endpoints (5.9%) despite measurable experimental bioactivity. Broadly, botanicals with stronger experimentally observed phenotypes tended to show activity across a larger number of predicted molecular and cellular endpoints, although this relationship was not uniform across extracts.

Several botanicals with experimentally observed cell health activity were also predicted to be active across multiple endocrine-disruption and genotoxicity assays. Goldenseal was predicted active in 53% of endocrine-related assays and 50% of genotoxicity assays, while aristolochia was predicted active in 47% and 83%, respectively. The genotoxicity predictions were also consistent with the relatively potent perturbation of nuclear morphology observed for these botanicals and their constituents (Figure 3.B). Together, these results generate specific hypotheses that endocrine- and genotoxicity-associated processes may contribute to the morphological responses observed for several bioactive botanicals.

## Discussion

In this study, we evaluated whether high-content phenotypic profiling could provide a scalable framework for characterising botanicals as complex mixtures. By screening botanical extracts alongside known constituents in primary human hepatocytes, we were able to compare extract- and constituent-level potency and identify compounds that may contribute to observed mixture activity. We further integrated Cell Painting profiles with a large reference atlas and ToxCast/Tox21 annotations to generate hypotheses about molecular and cellular processes associated with these phenotypes. Together, these analyses support the use of Cell Painting as a first-pass approach for prioritising active botanical extracts, nominating candidate constituent drivers, and guiding more targeted mechanistic follow-up.

We addressed the complex-mixture problem by profiling whole botanical extracts alongside a curated set of quantified constituents. This approach is complementary to other risk assessment complex-mixture methods characterisation. Effect-directed analysis (EDA) and toxicity identification evaluation (TIE) rely on fractionation, bioassay-guided tracking, and chemical identification to pinpoint specific toxicity (Escher et al., 2020). Our approach, centred on Cell Painting of individual constituent compounds alongside complete extracts, offers a fast, cost-effective means of characterising mixtures in their native, unfractionated form, preserving the whole-mixture context and potential constituent interactions that may be altered by fractionation. The trade-off is reduced mechanistic resolution: unlike EDA/TIE, Cell Painting alone makes it harder to fully resolve which individual or unknown constituents are associated with the observed activity. This limitation was partly addressed here by profiling extracts alongside quantified constituents, enabling extract-level phenotypes to be related to candidate constituent drivers.

Reference-atlas mapping provided an additional layer of interpretation beyond potency alone. Because the botanical and OASIS datasets were generated using closely matched Cell Painting protocols, their profiles could be aligned after batch correction and analysed within a common phenotypic space. Models trained on well-characterised OASIS compounds and associated ToxCast/Tox21 annotations were then applied to the botanical dataset to generate specific mode-of-action hypotheses. This strategy is consistent with a broader direction in computational toxicology and pharmacology, in which large reference datasets are used to relate molecular perturbations to cellular phenotypes and infer likely biological effects of less-characterised compounds (Meier et al., 2025; Trapotsi et al., 2022). Expanding such analyses with complementary morphology, transcriptomic, and bioactivity resources, including JUMP, L1000, ChEMBL, and Open TG-GATEs, could further improve the breadth and interpretability of these predictions [(Chandrasekaran et al., 2023; Gaulton et al., 2012; Igarashi et al., 2015; Subramanian et al., 2017)].

Several limitations should be considered when interpreting these findings. First, in vitro activity does not directly establish in vivo hepatotoxic risk. The concentrations reached in human liver after botanical consumption depend on absorption, distribution, metabolism, and excretion, and may differ substantially from nominal in vitro exposure concentrations [(Bianchi et al., 2024)]. Cell Painting bioactivity should therefore be interpreted as evidence of cellular perturbation rather than as a direct measure of hepatotoxicity, particularly because morphological responses may reflect adaptive or intended biological effects as well as adverse responses.

Second, interpretation of extract-level potency is complicated by mixture composition. A nominal extract concentration represents many constituents present at different abundances, and these abundances may vary between batches and products. Direct comparison between extract and pure-compound PODs therefore identifies plausible contributors but does not establish which compounds dominate the response in the intact mixture. Some relevant constituents may also have been underrepresented, absent from the tested panel, or present at concentrations that differ from those achieved in typical human exposure.

Third, the mechanistic predictions remain hypothesis-generating. Transfer performance for MT and LDH provided evidence that morphology-based models retained predictive signal when applied to the botanical dataset, but this does not directly validate predictions for endocrine, genotoxic, or other molecular endpoints. ToxCast/Tox21 annotations also capture only a subset of pathways relevant to hepatotoxicity and are themselves constrained by the biological fidelity of the underlying assay systems [(Xu et al., 2026; Ye et al., 2022)]. Predicted genotoxic, endocrine, and other pathway-level activities should therefore be considered priorities for orthogonal experimental validation rather than confirmed mechanisms.

Future studies could strengthen this framework by incorporating larger and more diverse Cell Painting reference atlases, additional target and pathway annotations, and complementary omics measurements. Integration with resources such as EU-OPENSCREEN and JUMP could broaden phenotypic coverage, while transcriptomic and biochemical reference datasets could improve mechanistic resolution [(Chandrasekaran et al., 2023; Wolff et al., 2025)]. Incorporating exposure information and direct quantification of constituent concentrations within botanical extracts would also help connect in vitro potency to realistic human exposures. Together, these extensions could improve the use of high-content phenotypic profiling for prioritising complex mixtures and directing targeted toxicological evaluation.

## Conclusion

In this study, Cell Painting provided sensitive concentration-dependent phenotypic readouts across botanical extracts and their constituents, while reference-atlas models enabled the generation of specific mechanistic hypotheses. Extract-level activity could often be linked to a subset of more potent constituents, with aristolochia and goldenseal among the most active botanicals profiled. These findings support morphological profiling combined with reference-based machine learning as a scalable framework for prioritising complex mixtures, nominating candidate constituent drivers, and guiding targeted follow-up studies.

## Acknowledgements

This work was supported in part by the Health and Environmental Sciences Institute’s (HESI Global) Botanical Safety Consortium. It is recognized via a Memorandum of Understanding between the US FDA, NIEHS, and HESI [MOU 225-19-032] that outlines joint commitments to a multisector and multidisciplinary Botanical Safety Consortium. This research was supported in part by the Intramural Research Program of the NIH, National Institute of Environmental Health Sciences (ZIA ES103381-05). This scientific initiative is primarily supported by in-kind contributions (from public and private sector participants) of time, expertise, and experimental effort. These contributions are supplemented by direct funding (that largely supports program infrastructure and management) that was provided by HESI Global’s corporate sponsors. A list of supporting organizations is available at https://botanicalsafetyconsortium.org/.

## Data and Code Availability

All original code and the proteomics data has been deposited at https://github.com/ewald-lab/2025_Ollitrault_BSC and is publicly available as of the date of publication.

Raw images have been deposited at https://broadinstitute.github.io/cellpainting-gallery and are publicly available as of the date of publication. They can be found at accession number ‘cpg0048-bsc’ in the “2024_09_20_BSC_Batch1” dataset folder.

Any additional information required to reanalyze the data reported in this paper is available from the corresponding author upon request.

