## Supplementary material for "Cell Painting and machine learning for mechanistic profiling of hepatotoxic botanicals and their constituents": [Supplementary] 20261020.pdf

### Supplementary data

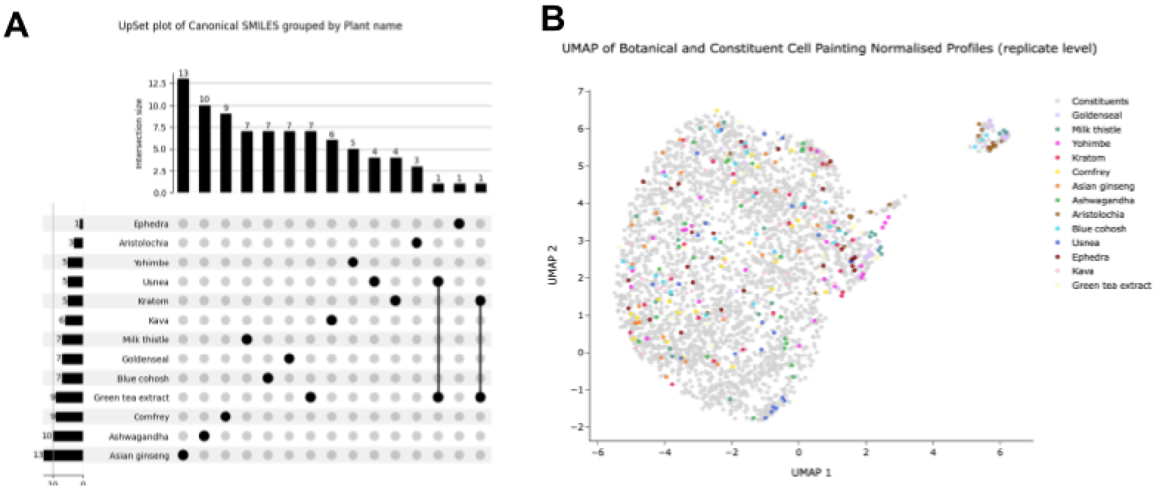


**Supplementary Figure 1. Constituents diversity of botanical extracts.**

A) Upset plot showing the number of unique and shared constituents tested across the botanical extracts.
B) UMAP of the phenotypic space of the botanicals and their constituents at the replicate level after normalisation

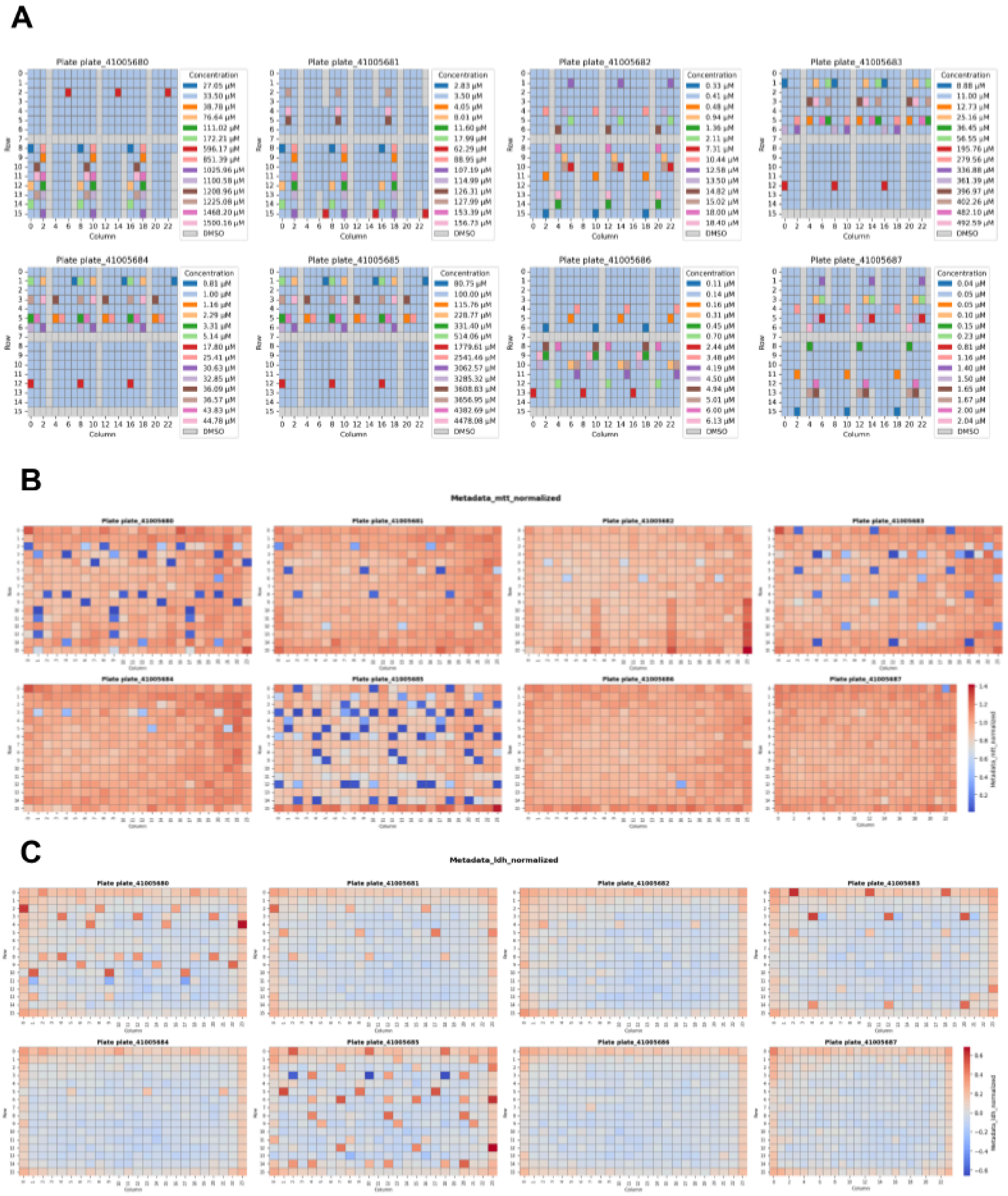


**Supplementary Figure 2. Experimental conditions and technical quality control.**

A) Distribution of botanical extract and constituent concentrations across wells and plates
B) Normalized RealTime-Glo™ MT luminescence values across wells and plates
C) Normalized CyQUANT LDH cytotoxicity values across wells and plates

**A**

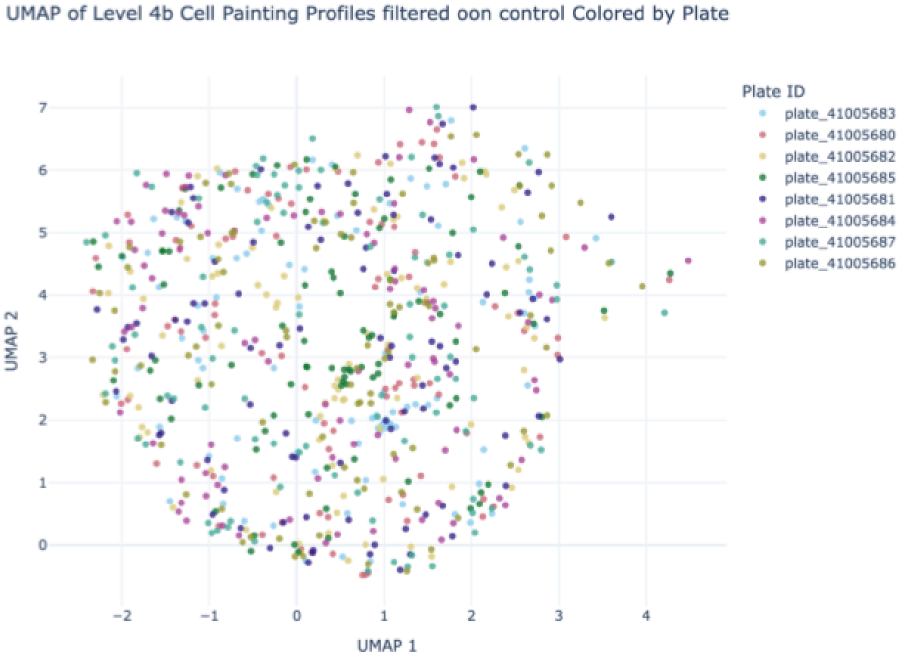


**B**

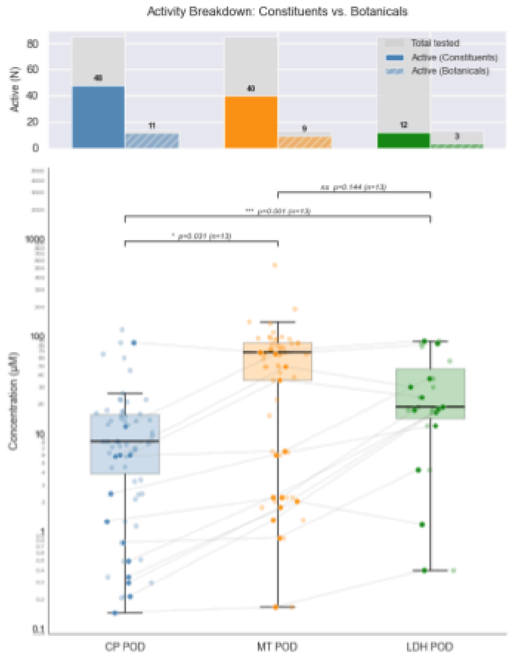


**Supplementary Figure 3. Bioactivity of botanical extracts and constituents**

A) UMAP of Level 4b Cell Painting phenotypic profiles after feature selection for DMSO control wells, colored by plate and replicate.
B) Bar plot showing the number of bioactive botanical extracts and constituents identified by Cell Painting, RealTime-Glo™ MT Luminescent Assay (MT), and CyQUANT LDH Cytotoxicity Assay (LDH). Comparison of points of departure (PODs) measured by Cell Painting, MT, and LDH. Botanical extracts and constituents active in all three assays are connected by grey lines across their assay-specific PODs. Pairwise Wilcoxon signed-rank tests performed on compounds common to all assays

are reported.

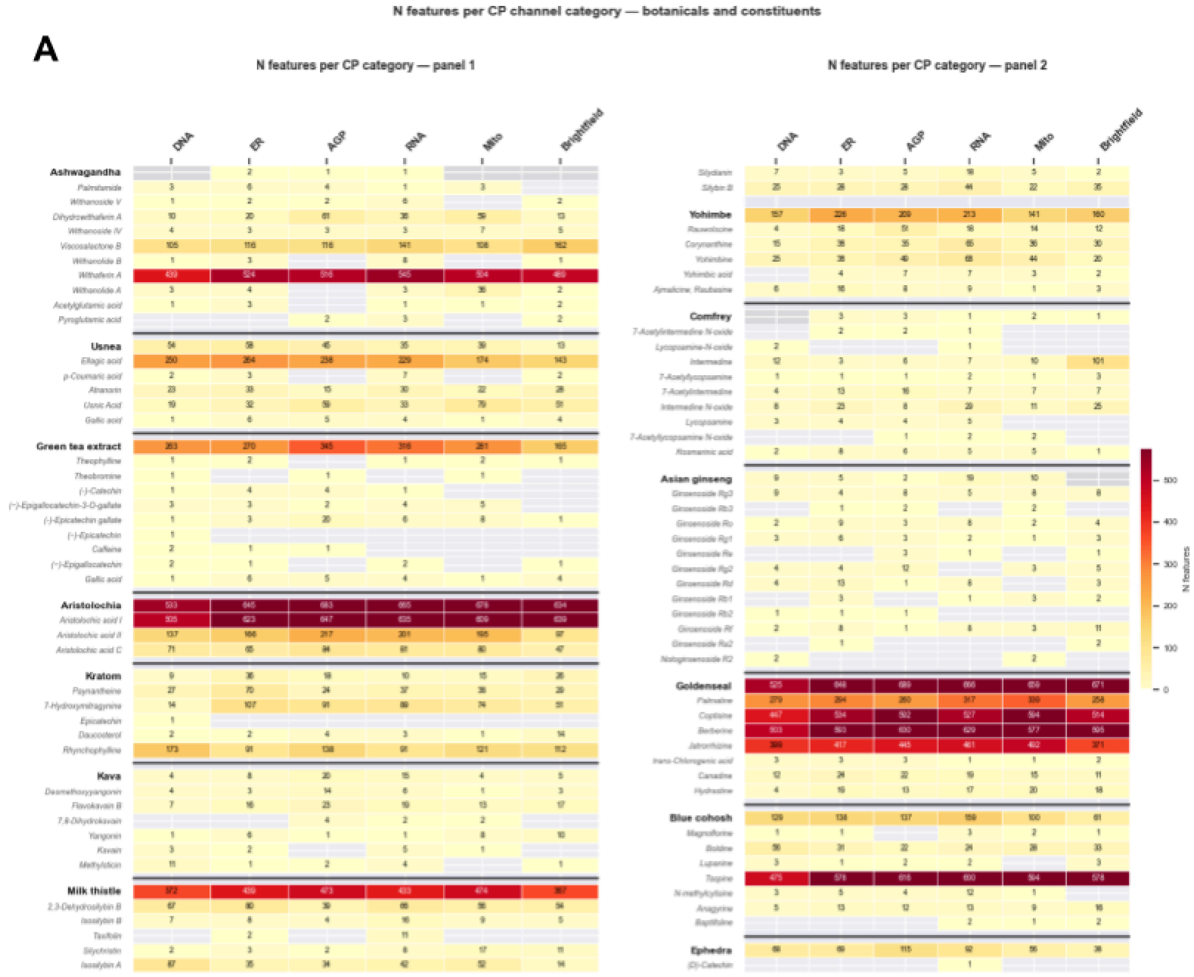


**Supplementary Figure 4. Phenotypic feature selection.**

A) Number of selected Cell Painting features for botanical extracts and constituents after feature filtering and interdependence score filtering of aggregated consensus phenotypic profiles (Level 6). Features are grouped by imaging channel

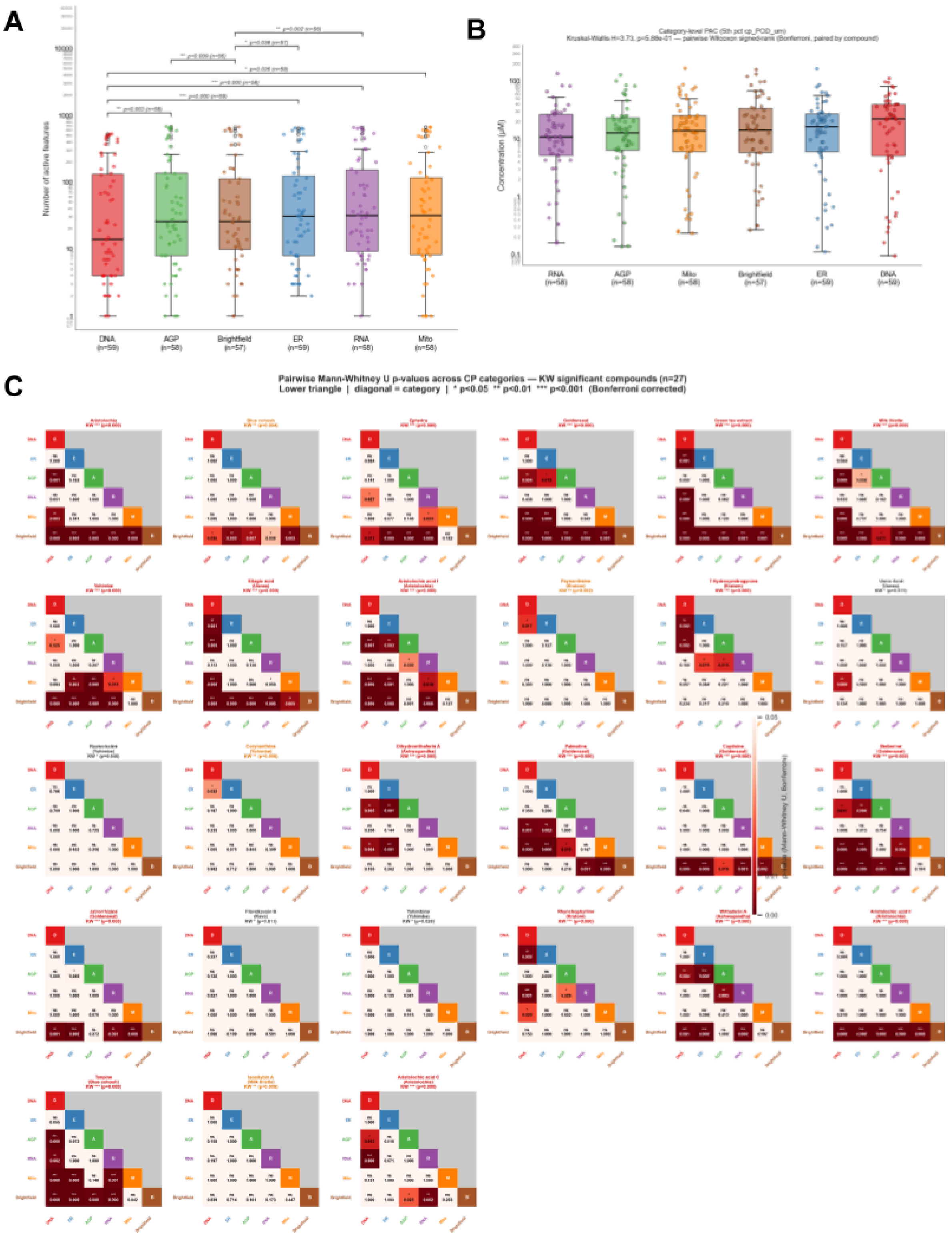


**Supplementary Figure 5. Channel-specific bioactivity.**

A) Number of active Cell Painting features grouped by imaging channel for active botanical extracts and constituents. Pairwise Wilcoxon signed-rank tests with Bonferroni correction are reported.

B) Channel-specific Cell Painting PODs for active botanical extracts and constituents. Pairwise Wilcoxon signed-rank tests with Bonferroni correction were not significant.

C) Pairwise Mann-Whitney U p-values comparing channel-specific POD distributions across botanical extracts and constituents

**A**

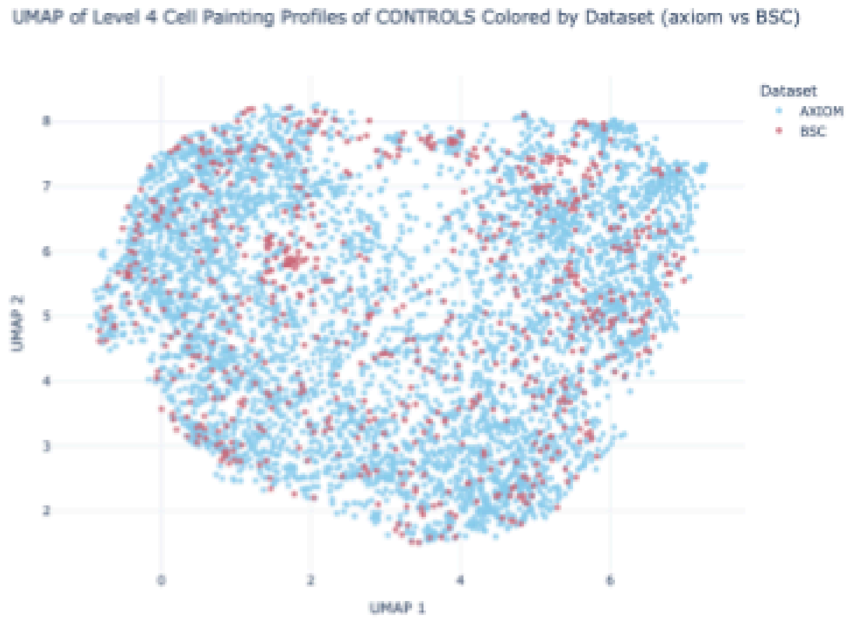


**Supplementary Figure 6. Phenotypic comparison of the OASIS and BSC datasets**

A) UMAP of Level 4 Cell Painting phenotypic profiles for control wells, colored by dataset.

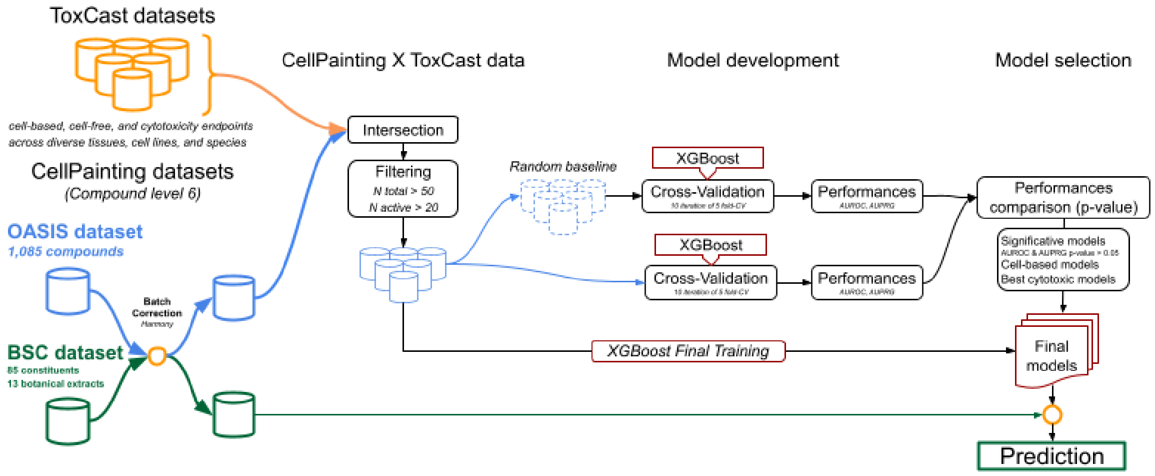


**Supplementary Figure 7. Prediction modeling workflow**

A) Schematic overview of the workflow used to develop Cell Painting-based predictive models and predict the mode of action of botanical extracts and their constituents.

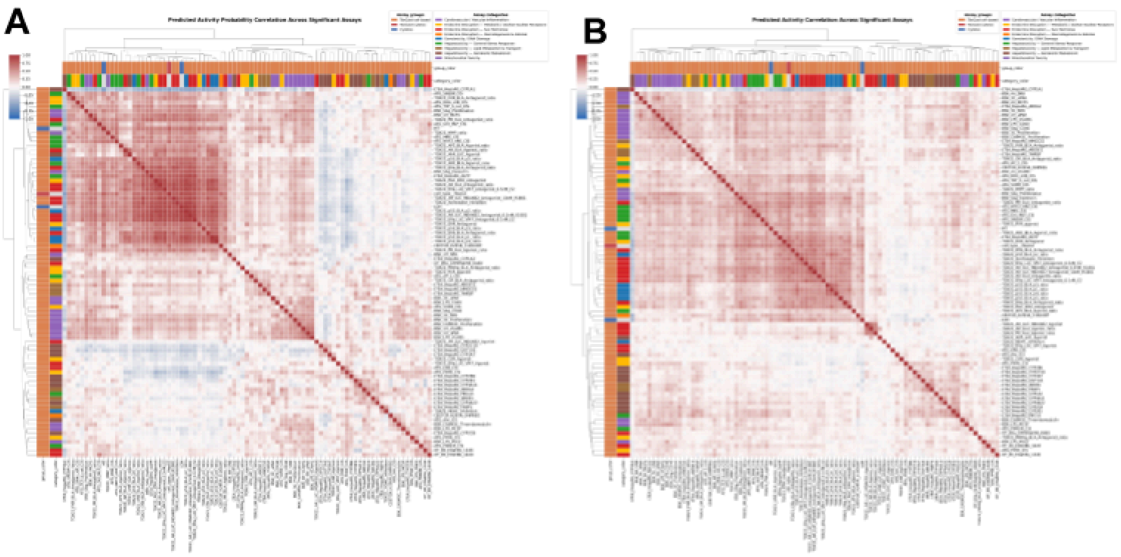


**Supplementary Figure 8. Evaluation of activity prediction models**

A) Clustermap of predicted activity probabilities for the botanical extract and constituent dataset across significant assays. Assays are annotated by biological group and category

B) Clustermap of predicted activity probabilities for the OASIS dataset across significant assays. Assays are annotated by biological group and category

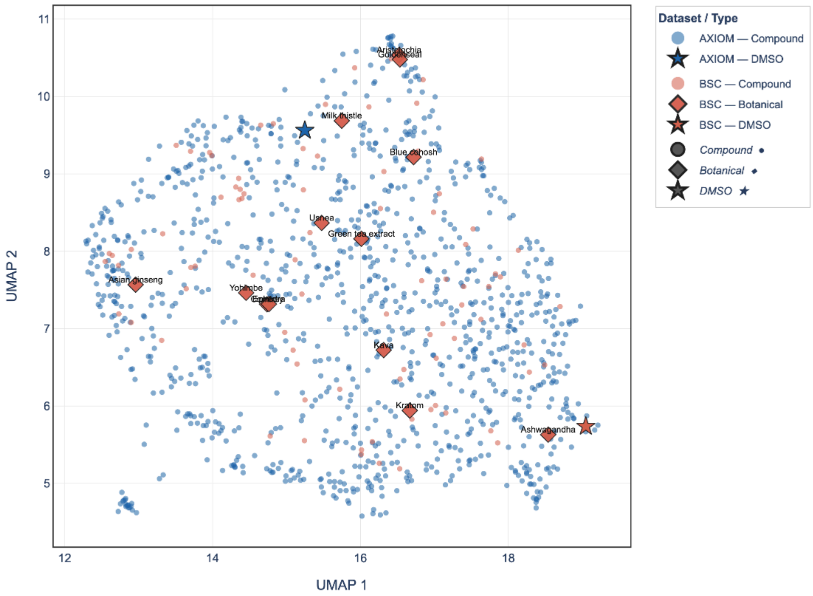


**Supplementary Figure 9.** UMAP of the profiles aggregated at the compound level. The dashed red circle corresponds to highly active cell health assays.

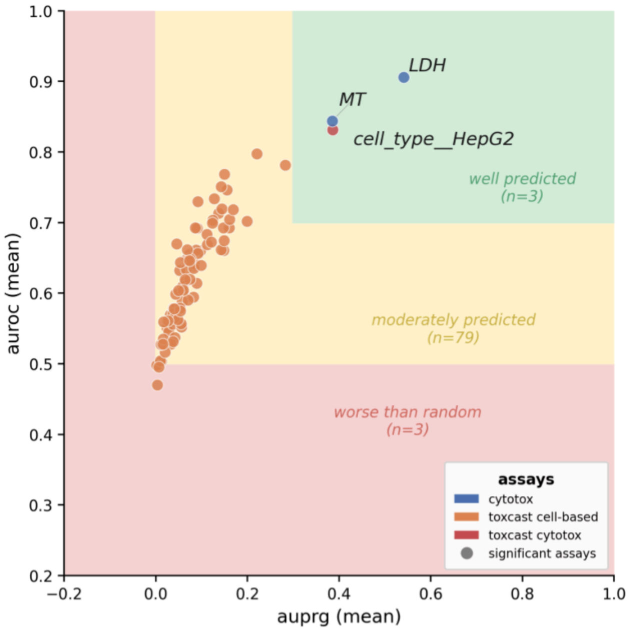


**Supplementary Figure 10.** Cross-validation performance of Cell Painting activity prediction models. Each point represents a model, plotted by mean AUROC and AUPRG. Filled circles indicate models that significantly outperformed a random baseline (paired t-test, $p < 0.05$); crosses indicate non-significant models.
